# Deep Learning-based Feature Extraction with MRI Data in Neuroimaging Genetics for Alzheimer’s Disease

**DOI:** 10.1101/2022.11.05.515286

**Authors:** Dipnil Chakraborty, Zhong Zhuang, Haoran Xue, Mark Fiecas, Xiaotong Shen, Alzheimer’s Disease Neuroimaging Initiative, Wei Pan

**Author notes:** Data used in preparation of this article were obtained from the Alzheimer’s Disease Neuroimaging Initiative (ADNI) database (adni.loni.usc.edu). As such, the investigators within the ADNI contributed to the design and implementation of ADNI and/or provided data but did not participate in analysis or writing of this report. A complete listing of ADNI investigators can be found at: http://adni.loni.usc.edu/wp-content/uploads/how_to_apply/ADNI_Acknowledgement_List.pdf.

## Abstract

The prognosis and treatment of the patients suffering from Alzheimer’s disease (AD) have been one of the most important and challenging problems over the last few decades. To better understand the mechanism of AD, it is of great interest to identify genetic variants associated with brain atrophy. Commonly in these analyses, neuroimaging features are extracted based on one of many possible brain atlases with FreeSurf and other popular softwares, which however may lose important information due to our incomplete knowledge about brain function embedded in these suboptimal atlases. To address the issue, we propose convolutional neural network (CNN) models applied to three-dimensional MRI data for the whole brain or multiple divided brain regions to perform completely data-driven and automatic feature extraction. These image-derived features are then used as endophenotypes in Genome-Wide Association Studies (GWAS) to identify associated genetic variants. When applied to the ADNI data, we identified several associated SNPs which have been previously shown to be related to several neurodegenerative/mental disorders such as AD, depression and schizophrenia. Code and supplementary materials are available at https://github.com/Dipnil07. The codes have been implemented using Python, R and Plink softwares.

## 1 Introduction

Alzheimer’s disease (AD) is an irreversible neurodegenerative disease that causes a decay in the brain tissue resulting in a loss of brain functions. It has been the most common cause of dementia, effecting roughly 5.8 million Americans aged 65 or above. AD has been identified as the sixth largest cause of death in US [1, 17]. The approximate cost of managing relevant finances of patients with Alzheimer’s disease was $277 billion in 2018 (Alzheimer’s Association, 2018) and is expected to almost double up by the next 20 years. There is no existing treatment to cure the disease. Unraveling the genetic architecture of AD may help to develop new preventive and treatment strategies.

The Alzheimer’s Disease Neuroimaging Initiative (ADNI) [28] is an ongoing research project on the development and progress of the disease. Magnetic resonance imaging (MRI) technologies are used to measure brain functions and structures and to assess the genetic architecture of the brain-related diseases using imaging-derived endophenotypes [7]. Data from these scans have been used to evaluate the neurological and psychriatic distortions created by diseases like Parkinson’s disease, Alzheimer’s disease, schizophrenia, autism and dementia. ADNI has provided a unique opportunity to combine data on MRI imaging and genetics.

A crucial challenge in this kind of study is to represent medical images into useful information that can ease clinical decision-making [13]. Traditionally, the most used approach has been image-on-scalar regression using mass univariate analysis (MUA). This method uses a general linear model at each voxel and calculates a voxel-wise test statistic. One drawback of the approach is that spatial correlation is not considered here which may result in low power. One of the potential remedies is to use all the voxels collectively as a tensor. Pan et al. (2019) [27] proposed a penalized approach to select subsets from the tensors after adjusting for the other covariates. In Miranda et al. [24], tensor partition regression modeling (TPRM) has been used to predict disease status by using structural medical resonance imaging (sMRI) data. Authors in Shi and Kang (2015) [35] model spatially varying functions as thresholded multiscale Gaussian processes from a bayesian point of view. In Feng et al. (2019) [9], authors developed a bayesian scalar-on-image regression model to combine the high dimensional image data and clinical data to predict different outcome states. The primary concern is that most of these statistical methods lose efficiency because of the high dimensionality of the array images as well as the complex structure. Advances in neuroimaging, deep learning, and genetics have opened new directions for feature extraction as endophenotypes to study the influence of genetics on brain structure and functions [32, 46].

A common approach to medical image analysis has been to use 2D convolutional neural networks (CNNs). While two dimensional CNNs have advantages in terms of data handling and limited GPU usage during optimization, the lack of knowledge about how to extract complete information using 2D patches prevents researchers from learning many important and robust information. Researches have shown that improved results are achievable when using the full volumetric data. In Zhao (2021) [48], the author used Bayesian semi-parametric model on ADNI data to jointly estimate voxel-specific heritability over whole brain imaging traits. In Bao (2021) [2], the author did an empirical study on the amyloid imaging and whole genome sequencing data using spatially connected voxels and showed higher estimated heritability measures. In this paper, we have explored the three-dimensional whole-brain structure MRI data obtained from ADNI.

In addition, for the genome-wide association analyses, researchers have extracted features from images using FreeSurf and other advanced normalization tools to use as phenotypes. In Grasby et al. (2020)[8], the authors used the surface area and cortical thickness of the cortex and cortical regions in a genome-wide association meta-analysis to link with several genetic variants. Zhao et al. (2020) [45] found significant genetic correlations between global functional connectivity measures and volumes of several brain regions. In Zhao et al.[47], the authors used consistent standard procedures via advanced normalization tools on MRI data and produced one hundred one region-based and overall brain volume phenotypes which were further used in GWAS analyses.

The work in this paper focuses on two important aspects: i) develop a completely data-driven feature extraction approach to ensure that the extracted features are related to AD and it is accomplished through CNN-based classification on the disease status, namely: Cognitively Normal (CN), Mild Cognitive Impairment (MCI), and Alzheimer’s disease (AD); ii) use these image-derived features as endophenotypes in a Genome-Wide Association Study (GWAS) to find out the genetic variants (SNPs) possibly related to AD. From the feature extraction point of view, it can be compared to the decomposed basis tensor layers mentioned in Tang et al. (2020) [40]. Sections 2 and 3 describe the data and methods used in the proposed deep learning and genome-wide association study. In Sections 4 and 5, contains the results of our study and relevant discussion, and Section 6 presents overall concluding remarks.

## 2 Data

In this paper, the imaging dataset as well as the genetic data were obtained from the Alzheimer’s Disease Neuroimaging Initiative (ADNI) website (http://adni.loni.usc.edu). ADNI was launched in 2003 by the National Institute of Biomedical Imaging and Bioengineering (NIBIB), the National Institute on Aging (NIA), thirteen pharmaceutical companies and two other agencies and was headed by Michael W. Weiner, MD, VA Medical Center and University of California, San Francisco. For most recent information and updates, see www.adni-info.org.

We first focus on the data containing the MRI scans of the 817 ADNI-1 subjects (188 early Alzheimer’s patients, 400 subjects showing Mild Cognitive Impairment, and 229 congnitively normal) and for the GWAS we consider the corresponding genetic data on 757 subjects. The ADNI data set contains three group of subjects, which are cognitively normal (CN) subjects, subjects with mild cognitive impairment (MCI) and subjects who have been diagnosed with Alzheimer’s disease (AD).

### 2.1 Preprocessing and description

In ADNI-1, the MRI data were obtained using different of MRI scanners with respect to particular protocols for each scanner. It included T1-weighted images which were obtained from volumetric three-dimensional sagittal MPRAGE or similar protocols using different resolutions [42]. The most commonly used image files were T1-weighted images obtained using 3D sagittal MPRAGE or other similar protocols. The magnetization-prepared rapid acquisition with gradient echo (MPRAGE) images from the LONI database have undergone the image processing steps including Gradwrap, B1 non-uniformity correction and N3 correction. Gradwrap corrects the image geometry distortion due to gradient non-linearity. B1 non-uniformity corrects for the intensity non-uniformity in the image. N3 is a histogram peak sharpening method which is applied to all images. Details on image preprocessing can be found at https://adni.loni.usc.edu/methods/mri-tool/mri-pre-processing/.

We used Brain Extraction tools (BET) [38] to extract the usable brain images. BET tool is a part of the well-known FMRIB (Functional Magnetic Resonance Imaging of Brain) Software Library (FSL) [43] containing tools for image and statistical analysis for functional, structural and diffusion MRI brain imaging data. Using BET, we deleted non-tissue regions from an image of the whole-brain structure (Figure 1, Left). In the whole image a large number of voxels are zero. This is expected as the brain is in a more centrally enclosed part of the image. Considering this fact and to minimize the computational cost we removed the encompassing regions with no information contained. Finally an image acquisition matrix of dimension 155 × 155 × 95 was used for the further studies.

**Figure 1:**
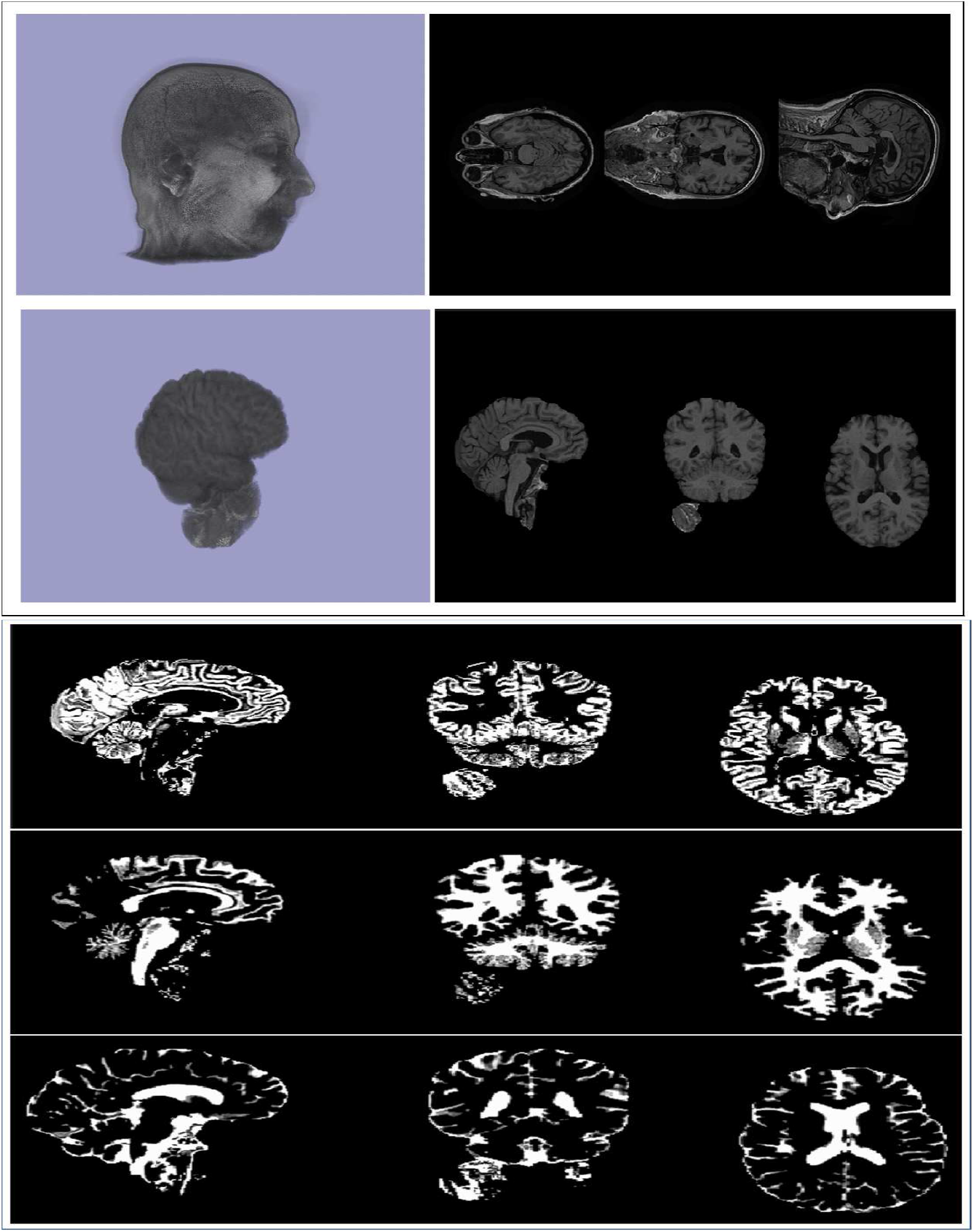
**Top-left:** An original MRI image and the corresponding axial views. **Bottom-left:** The extracted brain image using BET tool and its axial views. **Right:** Segmentation of the extracted brain image using FAST; the top to the bottom corresponds to the gray matter (GM), white matter (WM) and cerebrospinal fluid (CSF) images respectively.

## 3 Methods

### 3.1 Classification and feature extraction

Using deep learning to detect and classify diseases has gained a lot of attention in the recent years. Researchers, from across the globe, have been using convolutional neural networks (CNNs) for imaging as well as sequential [5] problems. Numerous studies in the last few years have focused on detecting Alzheimer’s disease and other types of dementias. Timely detection and diagnosis of Alzheimer’s disease helps to control or slow down the progression of the disease. Researchers have used the 3D CNN and its layers to understand the behavior of the disease spread.

In Kruthika et al.[19], auto encoders were used to extract features from the 3D patches showing a better result than the traditional 2D patches. In [22], author shows that the performance of the classification problem in case of dermoscopic analyses outperforms that of the raw images significantly. Researchers have also used LSTM (long short term memory) layers on top of three dimensional convolutional layers to improve the performance of the models. The 3D fully connected CNN layers were obtained for deep feature representations and LSTM was applied on these features to improve the performance. In Gao et al. [12], the authors improved the accuracy of AD, lesion and normal aging classification to 87.7% accuracy using a 7-layer deep three dimensional CNN on 285 volumetric CT head scans from Navy General hospital, China. In the literature of medical imaging, 3D CNN models were proposed for classification of 3D images as they could fully utilize the context information. However, development of 3D CNNs has been limited to simple architecture and small image size due to the high computational cost. In this section, we describe the 3D CNN architectures we used for the classification and feature extraction. We developed 3D CNN models with different architectures and different depths to classify AD patients into three categories, namely AD, MCI and CN, and extracted the output from a fully connected layer of 100 vectors known as neurons to be used as automatically extracted features. To further use the extracted features as endophenotypes in GWAS, we reduced the dimension of the feature vectors by Principal Components Analysis (PCA), then use top 10 PCs as endophenotypes.

#### 3.1.1 Whole-brain Structure

We started the study considering the extracted brain images obtained from BET tools. We used a 14-layered 3D convolutional neural network model (Figure 2) and added max pooling layers for dimension reduction and 3D convolution, as a natural adaption of 2D convolution. We also included batch normalization to each pooling layer. Throughout the CNN architecture, we used Rectified Linear Units (ReLU) [10] as the default activation function except a softmax [26] in the last layer for classification. In the proposed CNN model, we performed hyperparameter tuning to obtain the best possible results by modulating parameters such as: kernel size, steps, number of channels and dropout rate[21]. To minimize the overfitting issue we used two dropout layers with probability 30%.

**Figure 2:**
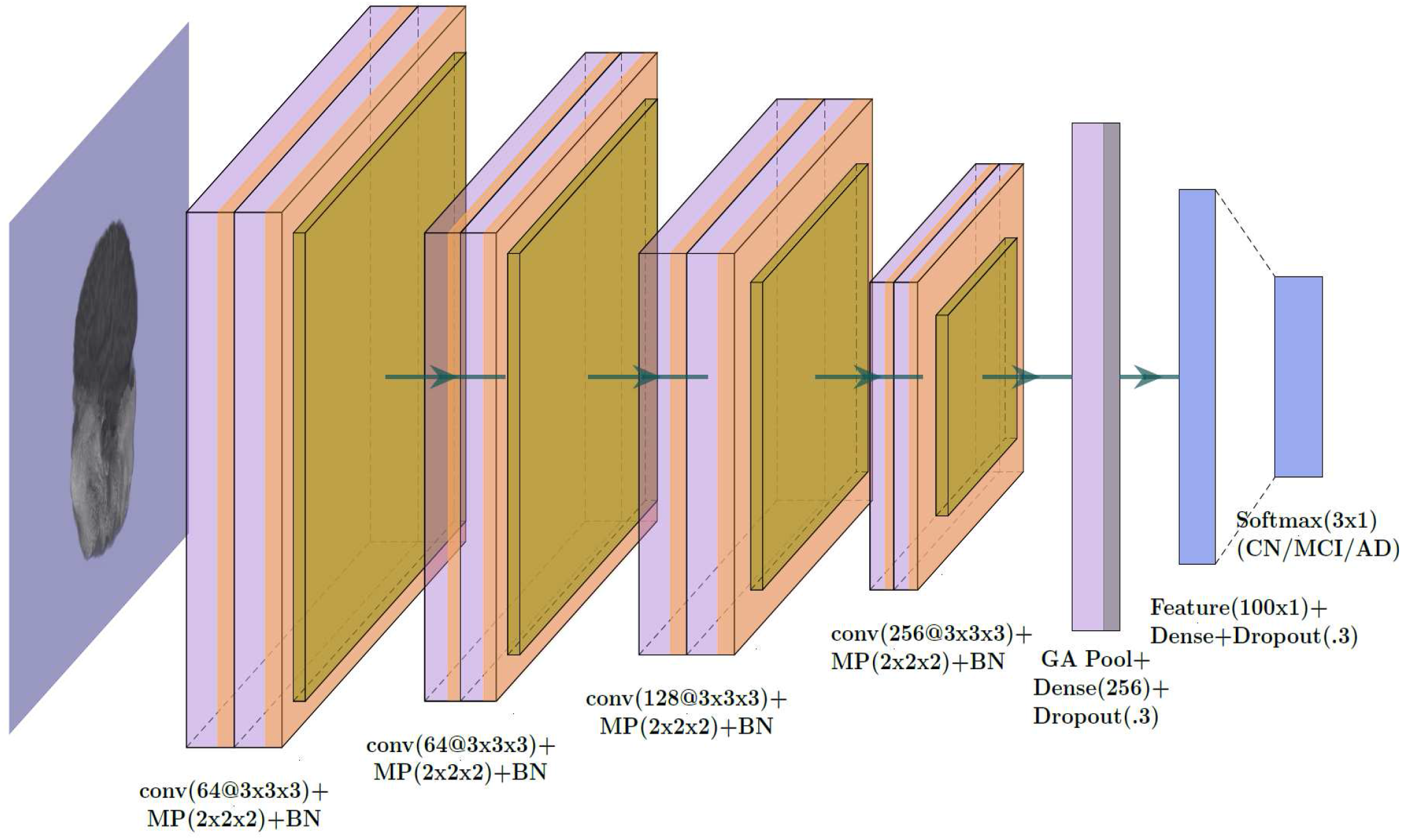
The Convolution Neural Network architecture used for the whole-brain structure. The penultimate dense layer of 100 neurons is extracted as the features.

From the CNN architecture, we extracted a fully connected layer with 100 neurons and used it as input corresponding to subjects for logistic regression and Support Vector Machine (SVM) classification models. Further we applied transfer learning algorithms. Transfer learning based neural network models have gained popularity in recent past. With some necessary modifications we implemented the idea of transfer learning models like AlexNet, ResNet50 and VGG16 in our study.

AlexNet [18], convolutional neural network was designed by Alex krizhevsky, in collaboration with Ilya Sutskever and Geoffrey Hinton to compete and win the first place at ImageNet Large Scale Visual Recognition Challenge (ILSVR) challenge. Visual Geometry Group (VGG16) [37] convolutional neural network proposed bu K. Simonyan and A. Zisserman secured second place in ILSVRC-2014 and makes an improvement over the AlexNet by using multiple 3 × 3 filters. Residual Network (ResNet) [14] based on a residual function won the first prize in ILSVRC-2015. We used the three dimensional counterpart of there models and applied transfer learning technique by replacing the top layer of the network with our classification layer.

To avoid over-fitting in our neural network architectures we opted for L2-norm regularization and implemented image augmentation by rotating the images at [-20,-10,-5,5,10,20] degrees respectively, i.e. for each input image in the training set now we will have six more images. In the model architectures, ReLU layers following convolution layers were used to introduce non-linearity necessary for class discrimination. The pooling layer were used to ensure that neurons don’t learn redundant information and the model does not grow too large while achieving some location-invariance. One of the primary concern of the study is to find out genetic variants associated with AD and hence, we decided to build a binary classification model as well based on only AD and CN subjects.

#### 3.1.2 Multibranch CNN

Using smaller parts of a 3D image often enables one to focus on locally most relevant information in classification. In our study, we divided the whole-brain structure along x, y and z axes in three parts respectively, resulting in 27 smaller and over-lapping 3D image patches. We trained a separate 14-layered CNN architecture with each of these pieces individually. The validation accuracies for these smaller patches were found to be noticeably higher for some parts than the test accuracies of the whole-brain CNN model. Thus we selected top five CNN models based on the accuracy levels and used them to build the final CNN model for the feature extraction (Figure 3). Similar to the whole-brain image scenario described earlier we collect a layer of fully connected neurons and used PCA to obtain top 10 PCs to be used further in the Genome-Wide Association Studies.

**Figure 3:**
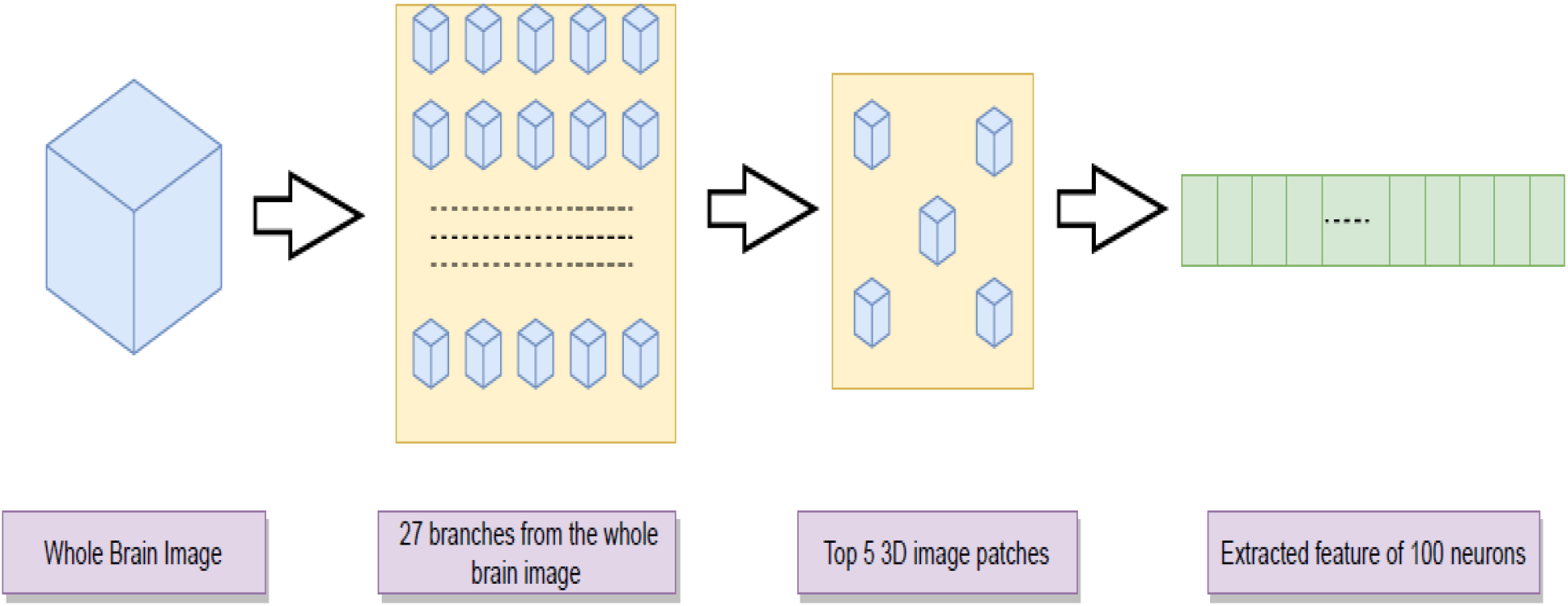
The pipeline for feature extraction using a multi-branch CNN model.

### 3.2 Segmentation

We applied the FMRIB’s Automated Segmentation Tool (FAST) [44] from the FSL software package to obtain individual 3D images corresponding to three tissue types, namely, grey matter (GM), white matter (WM) and cerebrospinal fluid (CSF) (Figure 1, Right). Further, we carried out CNN-based feature extraction and then GWAS on these segmented images similar to that discussed for whole-brain and multi-branch structures.

### 3.3 GWAS

One of the most important risk factors of AD is family history of dementia. Families with multiple members affected by AD are more susceptible to AD. Though we have made a significant progress in the research related to AD, perhaps most AD-related genetic variants and genes are still unknown. In this work, we applied PCA to choose the top 10 principal components (PCs) from the extracted feature vectors. The PCs were then each used as an endophenotype to conduct GWAS (while adjusting for additional covariates such as age, sex, education, handedness etc).

### 3.4 PC vs Grey Matter Volume of ROIs

A number of recent studies have shifted their focus to develop machine learning models that can be interpreted easily. In Ribeiro et al. [30], the authors discuss about using explainable methods to be able to trust the predictions made by the “black-box” models like CNNs. In this work, we used linear regression to investigate the relations of the extracted principal components with some commonly used ROIs as already available in the ADNI website, such as the GM volumes of Left Hippocampus (HIPPL), Para Left Hippocampus (PARAHIPPL) and CEREBELLUM etc, extracted from 1.5T ADNI MPRAGE MRI segmented gray matter maps using longitudinal VBM (voxel based morphometry). These ROIs have been previously shown to change significantly between the AD and control subjects [23].

### 3.5 Evaluation Criteria

The receiver operating characteristic (ROC) [4] curve is used by plotting the True Positive Rate (TPR) against the False Positive Rate (FPR). The area under ROC curve (AUC) is one of the most widely used metrics for classification. The training AUCs reported in the paper are based on the validation sets. We have used the AUCs obtained from the models to compare their prediction accuracy. We split the whole image dataset into training (80%), validation (10%) and test (10%) subsets and calculated the (test) AUC with the test subset; then the above process was repeated for 10 times and we calculated the average (test) AUCs reported in this paper. The models with higher prediction accuracies were used for the GWAS.

## 4 Results

### 4.1 Whole-brain Structure

The training AUC for the CNN model corresponding to the images of the whole-brain structure is 0.88 and the test AUC is 0.72. Performing GWAS with the top 10 PCs each as the phenotype yielded 149 SNPs at p-value *≤* 5× 10^*−*6^, including rs2075650, which has been found to be consistently associated with AD [29, 6]. Apolipoprotein E (APOE) gene and the TOMM40-APOE locus, tagged by rs2075650 in the translocase of outer mitochondrial membrane 40 (TOMM40) gene are both considered to be responsible for AD. TOMM40 rs2075650 (chromosome 19q13.32) has been one of the most important SNPs found to be associated with AD. TOMM40 encodes translocase of the mitochondrial outer membrane (TOM) complex through mitochondrial channel protein TOM40 [15]. Changes in the mitochondrial metabolism have been accepted as a feature in patients of Alzheimer’s Disease. eMERGE network showed an association of rs2075650 with AD using electronic medical records [6].

Next, we implemented logistic regression considering the extracted feature vectors as inputs for corresponding subjects and the model produced a training AUC of 0.78 and test AUC 0.59. Similarly, a support vector machine (SVM) yielded a training and test AUC of 0.79 and 0.61 respectively. CNN models based on transfer learning, such as AlexNet, ResNet50, VGG16, produced training AUC within the range 0.84-0.87 and test AUC of 0.69-0.72. None of these models provided improved accurarcy measures in the classification problem and hence were dropped in future consideration for feature extraction and GWAS.

Over-fitting and small sample sizes are two of the main growing concerns of neural network models, especially in medical image data analyses. We trained the CNN model with augmentation by rotating the images in six different angles. Introducing augmentation of the training sample made the input richer with information and improved the AUC measure of the test dataset up to 0.75. GWAS based on the derived features from this model yielded 3 SNPs at p-values *≤* 5 × 10^*−*8^, including rs2075650, rs11580593 and rs823955, and 35 SNPs at the marginal significance level of 5 × 10^*−*6^.

Many researches have been conducted with the primary focus to identify the genetic differences between AD and CN subjects. For that purpose, we used a CNN model to address this binary classification problem which yielded a training AUC of 0.96 and test AUC of 0.90. A genome-wide association scan produced 2 SNPs at the p-value significance level of 5× 10^*−*8^ and 53 SNPs at 5× 10^*−*6^ with the inclusion of rs7836628 (4.21 × 10^*−*8^) and rs12708438 (4.33 × 10^*−*8^) SNPs. This list of SNPs also include rs2196315 [chr8:130787049] (8.02× 10^*−*8^) corresponding to gene ADCY8 (Intron Variant) which has been found to be connected with Dissociative Amnesia (corresponding to rs726411 located in 8q24.22 region [25] and rs263238 [11] with *p* = 2.40× 10^*−*6^), whereas SNP rs2395095 of Adenosine Kinase (ADK) gene has been previously found to be responsible to trigger cognitive impairment, and seizures [3, 31]. The Manhattan plot on the top-left panel of Figure 4 corresponding to the AD vs CN classification shows a large number of significant SNPs on chromosome 15 (chr15:29176591, chr15:29175402, chr15:29190239, chr15:29225416), which has been an increasing focus in the field on neurology and has shown association with several neurodegenarative/mental/brain diseases such as Dyslexia, Autism, Ring 15 Chromosome syndrome, Angelman syndrome and Hyperlexia [36]. Table 1, summarizes the most significant SNPs corresponding to the principal components obtained from the discussed models along with their chromosomal positions and p-values from the relevant GWAS. We used the CNN model discussed in the whole-brain section individually for GM, WM and CSF segments. The training and testing AUC for the CNN models in case of GM images were found to be 0.82 and 0.70 respectively. Similarly, the training and testing measures of AUC for WM images were 0.88 and 0.68 and those for the CSF images were 0.77 and 0.54 respectively. The relevant GWAS did not identify any significant SNPs at p-value 5 × 10^*−*8^.

**Table 1:** Models based on whole-brain images with contributing principal component and significant SNPs with their chromosomal positions and p-values.

| Models | Test AUC | PC involved | SNP | Chromosome | P-value |
| --- | --- | --- | --- | --- | --- |
| Augmentation with respect to whole-brain | 0.75 | 3 | *rs1158059* | 1 | $2.06 \times 10^{-9}$ |
| | | 2 | *rs2075650* | 19 | $4.58 \times 10^{-8}$ |
| | | 4 | *rs823955* | 3 | $4.72 \times 10^{-8}$ |
| AD vs CN Binary Classification | 0.90 | 4 | *rs7836628* | 8 | $4.21 \times 10^{-8}$ |
| | | 7 | *rs12708438* | 15 | $4.33 \times 10^{-8}$ |

**Figure 4:**
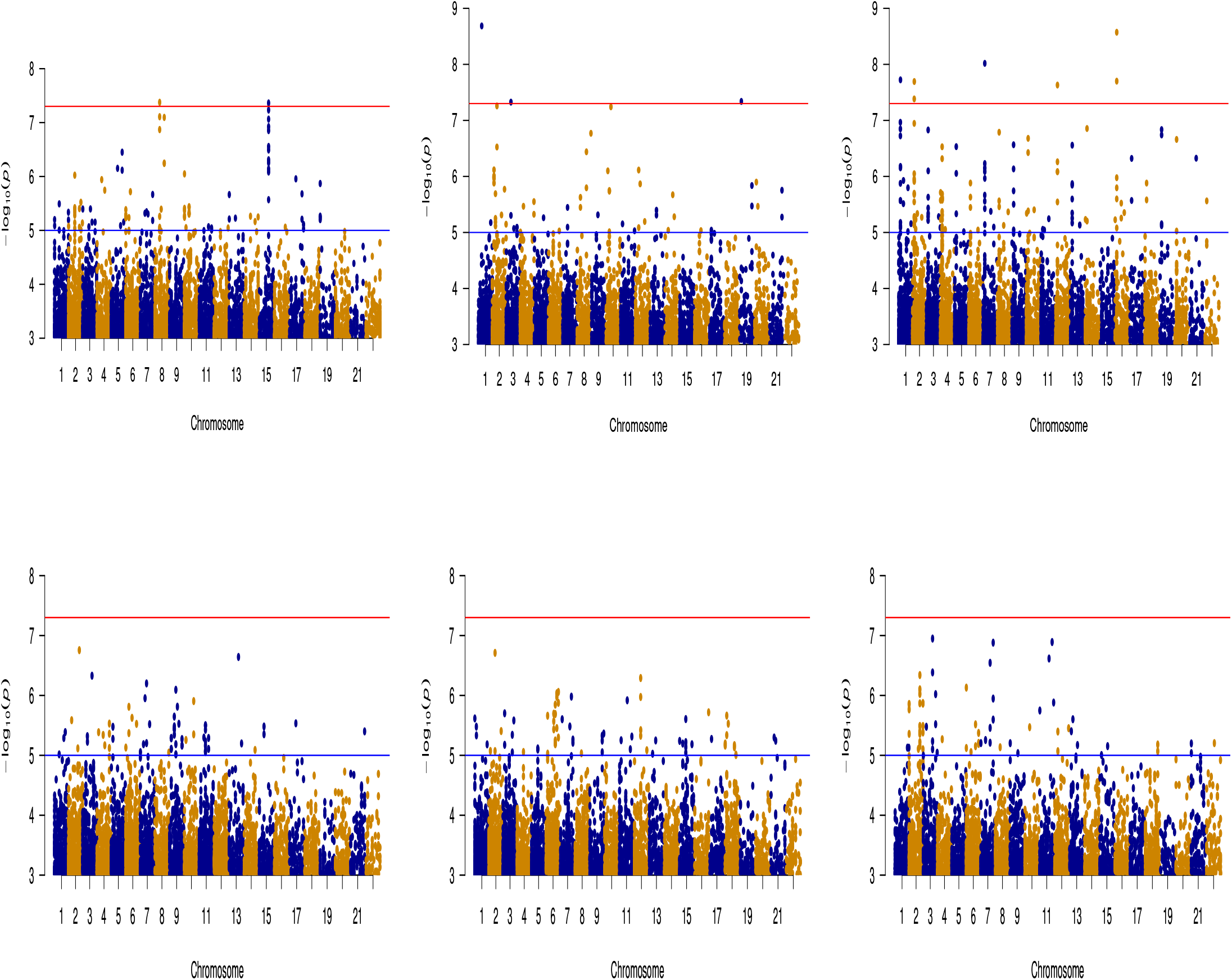
The Manhattan Plots. **Top panels:** The left one corresponds to binary classification of AD vs CN, the one in the middle is for whole-brain structure with augmentation, and the right one is for the multi-branch model for the whole-brain structure. **Bottom panels:** The left one is for the multi-branch model on GM images, middle one is for the multi-branch model on WM images, and the right one is for the multi-branch model on CSF images.

### 4.2 Multi-branch CNN

For the multi-branch CNN, we trained a separate CNN model for each of the 27 sub 3D image patches. We selected the top 5 patches and combined them together to obtain our final model. The training accuracy obtained for the final model was 0.86 and the testing accuracy thus obtained was 0.76. The extracted feature vector from the final model was to obtain the top 10 Principal components. The GWAS conducted with these PCs identified 8 SNPs at p-value 5 × 10^*−*8^ and 87 SNPs at p-value 5 × 10^*−*6^. The list of significant SNP includes rs1397645 [ch4:121036579] from Neuron Derived Neurotrophic Factor (NDNF) which helps to protect and culture hippocamal neurons against amyloid beta-peptide toxicity[20]. Next, we used the multibranch model technique on GM images. The training and test AUC measures obtained were 0.84 and 0.74. With the GWAS study, we identified that 35 SNPs are significant at p-value 5 × 10^*−*6^. This list of SNPs includes rs173754 (3.33× 10^*−*6^) which has been shown responsible for ADHD (Attention Deficit Hyperactivity Disorder).

This list of SNPs includes rs9257694 [chr6:29306709] (1.55 × 10^*−*6^) associated with OR14J1 (olfactory receptor family 14 subfamily J member 1). Though no concrete conclusions have been drawn, but abnormalities and impaired functions of the olfactory system have been reported in AD, and the these symptoms have been seen to appear earlier than many other symptoms. Table 2, gives us a glimpse of the most significant SNPs corresponding to the principal components obtained from the multibranch models along with their chromosomal position and calculated P-values from the relevant GWAS studies. Next, we used the multibranch model technique on GM images. The training and test AUC measures obtained were 0.84 and 0.74. With the GWAS study, we identified that 35 SNPs are significant at p-value 5 × 10^*−*6^. This list of SNPs includes rs173754 (3.33 × 10^*−*6^) which has been seen responsible for ADHD (Attention Deficit Hyperactivity Disorder).

**Table 2:** Multi-branch models with contributing principal components and significant SNPs with their chromosomal positions and p-values.

| Models | Test AUC | PC involved | SNP | Chromosome | P-value |
| --- | --- | --- | --- | --- | --- |
| Multi-branch | 0.76 | 4 | *rs1397645* | 8 | $9.40 \times 10^{-8}$ |
| | | 7 | *rs10514441* | 16 | $2.67 \times 10^{-9}$ |
| | | 9 | *rs17148126* | 7 | $9.59 \times 10^{-9}$ |
| Multi-branch on GM images | 0.74 | 2 | *rs9257694* | 6 | $1.55 \times 10^{-6}$ |
| | | 3 | *rs173754* | 5 | $3.33 \times 10^{-6}$ |

The multi-branch model on the WM images produced AUC measures of 0.82 and 0.70 on the training and test datasets respectively. GWAS based on this model identified 35 SNPs (Figure 4: Bottom panel on the left) at the marginal significance level of 5 × 10^*−*6^. On the other hand, the model on the CSF images produced AUC of 0.77 on training data and 0.66 on the test data and the further GWAS analysis yielded 21 SNPs, but the SNPs were not found to be associated with any well known genes. The list of significant SNPs include rs11710427, rs15820, rs9791663, rs9791663, rs3132030, rs10880942 etc. The bottom panels of Figure 4 show the Manhattan plots for all the segment-wise multibranch models.

### 4.3 PC vs Grey Matter Volumes of ROIs

To enhance the interpretation of the extracted features, we investigated the relation of the mean grey matter volmes of brain ROIs and principal components using linear regression. Figure 5 shows for the 2nd and 3rd principal components obtained from the whole-brain structure with augmentation. The x-axis shows the ROI indices and the y-axis shows the *−* log(p value) obtained from the regression. The results demonstrate that mean grey matter volumes of the ROIs such as left/right hippocampus (HIPPL/R), right parietal superior (PARIETSUPR), and several parts of cerebellum (left/right hemispheric lobule VIIb, CEREB7BL/R; left/right hemispheric lobule VIII, CEREB8L/R) show a higher level of significance than the other regions. However, using the principal components we found SNPs which were not identified using the ROIs only. One of the reasons may be that apart from the major effect from one brain ROI, the principal components may have information from other brain regions as well. For instance, the 3rd PC of the augmented model was found associated with *rs*1158059 from gene SNAP91 which has also been found to be involved in AD pathways [41]; the 7th principal component of the multi-branch model with the whole brain method identifies SNP *rs*10514441 related to gene WWOX which plays an important role AD through interactions with its protein partners and cell pathology and degeneration [39]; These brain regions [33, 16] have been previously shown to be associated with AD. Moreover, the SNP *rs*2075650 associated with the 2nd principal component (Table 1) has been shown to be strongly associated with hippocampus volume of the brain [34, 29]. The relations among the principal components and the available grey matter volumes of the ROIs provide a credence to the used models and enhance the interpretability of the models. See supplementary material for more plots.

**Figure 5:**
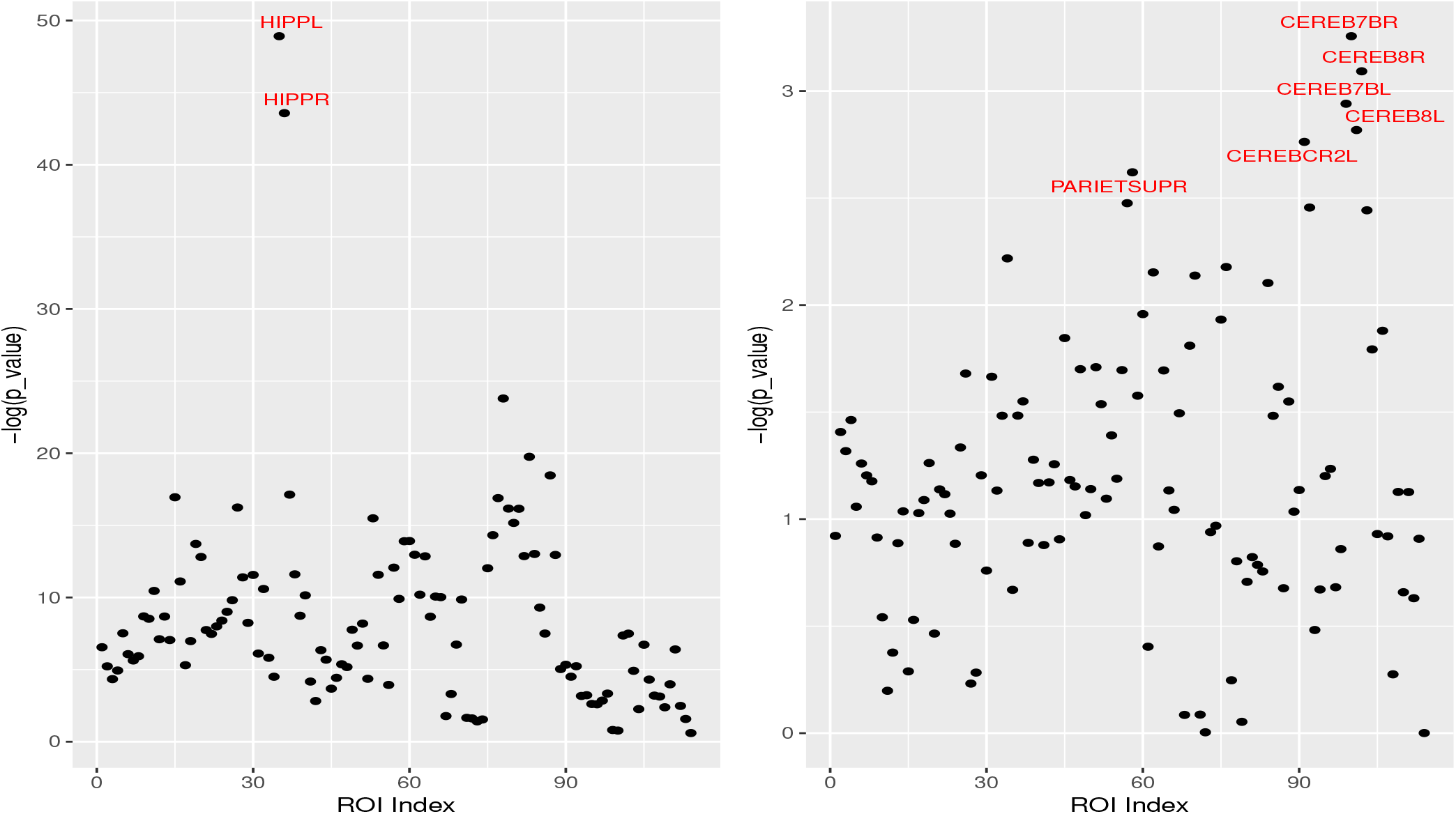
Marginal associations between each of the brain ROIs and one principal component. X-axis shows the different ROI indices and Y-axis shows the –log(p value) obtained from the regression. The left panel is for PC2 and the right one is for PC3, both obtained from the whole-brain augmentation.

## 5 Discussion

In this paper, we have explored completely data-driven and automated feature extraction and using them as phenotypes in the corresponding GWAS primarily based on two techniques: the first corresponding to the CNN models trained on the whole images, and the second to multi-branch CNN models trained on divided sub-parts of the whole images. We noticed that the initial CNN model based on the whole-brain structure was not the best performing model in terms of the prediction accuracy. The follow-up GWAS for the model also failed to capture significant SNPs at lower p-values cut-offs. One problem that the model suffered from, was over-fitting. To tackle the issue, we implemented an augmentation method and performed the same steps on the augmented set of images, producing much more accurate results along with significant SNPs (Table 1). Next, we adopted the binary classification problem between the AD and CN subjects, instead of the existing tertiary classification problem. The used model, as expected, performed better reaching up to 0.90 AUC measure on the test dataset. The relevant GWAS brought out significant SNPs at the usual genome-wide significance levle of p-value *≤* 5 × 10^*−*8^ (Table 1).

The most important contribution of the paper is perhaps the idea of the multi-branch CNN models where we trained the convolutional neural network models individually on 27 sub-parts of each whole-brain image and identified top five patches with the highest validation accuracies to be included in the final model. Focusing only on the pertinent patches of the images helps us not only localize relevant brain regions, but also retrieve maximum local information, thus boosting the AUC measures of the models. Features pulled out from these models also pointed out more significant SNPs compared to their whole-image counterparts (Table 2).

## 6 Conclusion

Alzheimer’s disease is the most common form of dementia. The worldwide number of the patients suffering from this disease are increasing everyday. The disease starts to develop years before the first noticeable symptoms start to appear. The study in this paper focuses on two major issues related to AD. Firstly, it proposes a method of automated feature extraction, compared to focusing on some pre-defined ROIs using existing software, through classification of AD patients. The study shows that using multi-branch CNN models, instead of that for the regular whole-brain structures, provides an increase in the prediction accuracy and hence should be considered as an option for future brain related researches. Secondly, the study identified the genetic variants related to AD. The work links the associated SNPs from the available genetic data to the imaging data of the brain. Some of the SNPs identified in our work have been previously shown to be related to mental/brain disorders, including schizophrenia, depression and dementia.

## Data Availability

The imaging dataset as well as the genetic data used in this work can be obtained from the Alzheimer’s Disease Neuroimaging Initiative (ADNI) website (http://adni.loni.usc.edu)

## Acknowledgments

This research was funded by NIH grants R01 AG069895, RF1 AG067924, R01 AG065636, R21AG057038 and by the Minnesota Supercomputing Institute at the University of Minnesota.

Data collection and sharing for this project was funded by the Alzheimer’s Disease Neuroimaging Initiative (ADNI) (National Institutes of Health Grant U01 AG024904) and DOD ADNI (Department of Defense award number W81XWH-12-2-0012). ADNI is funded by the National Institute on Aging, the National Institute of Biomedical Imaging and Bioengineering, and through generous contributions from the following: AbbVie, Alzheimer’s Association; Alzheimer’s Drug Discovery Foundation; Araclon Biotech; BioClinica, Inc.; Biogen; Bristol-Myers Squibb Company; CereSpir, Inc.; Cogstate; Eisai Inc.; Elan Pharmaceuticals, Inc.; Eli Lilly and Company; EuroImmun; F. Hoffmann -La Roche Ltd and its affiliated company Genentech, Inc.; Fujirebio; GE Healthcare; IXICO Ltd.; Janssen Alzheimer Immunotherapy Research & Development, LLC.; Johnson & Johnson Pharmaceutical Research & Development LLC.; Lumosity; Lundbeck; Merck & Co., Inc.; Meso Scale Diagnostics, LLC.; NeuroRx Research; Neurotrack Technologies; Novartis Pharmaceuticals Corporation; Pfizer Inc.; Piramal Imaging; Servier; Takeda Pharmaceutical Company; and Transition Therapeutics. The Canadian Institutes of Health Research is providing funds to support ADNI clinical sites in Canada. Private sector contributions are facilitated by the Foundation for the National Institutes of Health (www.fnih.org). The grantee organization is the Northern California Institute for Research and Education, and the study is coordinated by the Alzheimer’s Therapeutic Research Institute at the University of Southern California. ADNI data are disseminated by the Laboratory for Neuro Imaging at the University of Southern California.

## References

[1] 2018 alzheimer’s disease facts and figures. Alzheimer’s & Dementia, 14(3):367–429, mar 2018.

[2] Jingxuan Bao, Zixuan Wen, Mansu Kim, Xiwen Zhao, Brian N. Lee, Sang-Hyuk Jung, Christos Davatzikos, Andrew J. Saykin, Paul M. Thompson, Dokyoon Kim, Yize Zhao, and Li Shen. Identifying highly heritable brain amyloid phenotypes through mining alzheimer’s imaging and sequencing biobank data. In Biocomputing 2022. WORLD SCIENTIFIC, nov 2021.

[3] Detlev Boison. Adenosine kinase: Exploitation for therapeutic gain. Pharmacological Reviews, 65(3):906–943, apr 2013.

[4] Andrew P. Bradley. The use of the area under the ROC curve in the evaluation of machine learning algorithms. Pattern Recognition, 30(7):1145–1159, jul 1997.

[5] Dipnil Chakraborty and Akash Roy. Time series methodology in STORJ token prediction. In 2019 International Conference on Data Mining Workshops (ICDMW). IEEE, nov 2019.

[6] Joshua C Denny, Lisa Bastarache, Marylyn D Ritchie, Robert J Carroll, Raquel Zink, Jonathan D Mosley, Julie R Field, Jill M Pulley, Andrea H Ramirez, Erica Bowton, Melissa A Basford, David S Carrell, Peggy L Peissig, Abel N Kho, Jennifer A Pacheco, Luke V Rasmussen, David R Crosslin, Paul K Crane, Jyotishman Pathak, Suzette J Bielinski, Sarah A Pendergrass, Hua Xu, Lucia A Hindorff, Rongling Li, Teri A Manolio, Christopher G Chute, Rex L Chisholm, Eric B Larson, Gail P Jarvik, Murray H Brilliant, Catherine A McCarty, Iftikhar J Kullo, Jonathan L Haines, Dana C Crawford, Daniel R Masys, and Dan M Roden. Systematic comparison of phenome-wide association study of electronic medical record data and genome-wide association study data. Nature Biotechnology, 31(12):1102–1111, ec 2013.

[7] Lloyd T. Elliott, Kevin Sharp, and ….. Stephen M. Smith. Genome-wide association studies of brain imaging phenotypes in UK biobank. Nature, 562(7726):210–216, oct 2018.

[8] Katrina L. Grasby et al. The genetic architecture of the human cerebral cortex. Science, 367(6484):eaay6690, 2020.

[9] Xiangnan Feng, Tengfei Li, Xinyuan Song, and Hongtu Zhu. Bayesian scalar on image regression with nonignorable nonresponse. Journal of the American Statistical Association, 115(532):1574–1597, 2020. PMID: 33627920.

[10] Kunihiko Fukushima. Neocognitron: A self-organizing neural network model for a mechanism of pattern recognition unaffected by shift in position. Biological Cybernetics, 36(4):193–202, apr 1980.

[11] S J Furney,, A Simmons, G Breen, I Pedroso, K Lunnon, P Proitsi, A Hodges, J Powell, L-O Wahlund, I Kloszewska, P Mecocci, H Soininen, M Tsolaki, B Vellas, C Spenger, M Lathrop, L Shen, S Kim, A J Saykin, M W Weiner, and S Lovestone. Genome-wide association with MRI atrophy measures as a quantitative trait locus for alzheimer’s disease. Molecular Psychiatry, 16(11):1130–1138, nov 2010.

[12] Xiaohong W. Gao, Rui Hui, and Zengmin Tian. Classification of CT brain images based on deep learning networks. Computer Methods and Programs in Biomedicine, 138:49–56, jan 2017.

[13] Robert J. Gillies, Paul E. Kinahan, and Hedvig Hricak. Radiomics: Images are more than pictures, they are data. Radiology, 278(2):563–577, feb 2016.

[14] Kaiming He, Xiangyu Zhang, Shaoqing Ren, and Jian Sun. Deep residual learning for image recognition. In 2016 IEEE Conference on Computer Vision and Pattern Recognition (CVPR), pages 770–778, 2016.

[15] Nicholas J Hoogenraad, Linda A Ward, and Michael T Ryan. Import and assembly of proteins into mitochondria of mammalian cells. Biochimica et Biophysica Acta (BBA) - Molecular Cell Research, 1592(1):97–105, sep 2002.

[16] Heidi I L Jacobs, David A Hopkins, Helen C Mayrhofer, Emiliano Bruner, Fred W van Leeuwen, Wijnand Raaijmakers, and Jeremy D Schmahmann. The cerebellum in alzheimer’s disease: evaluating its role in cognitive decline. Brain, 141(1):37–47, jul 2017.

[17] Ellen A Kramarow and Betzaida Tejada-Vera. Dementia mortality in the united states, 2000-2017. National vital statistics reports : from the Centers for Disease Control and Prevention, National Center for Health Statistics, National Vital Statistics System, 68:1–29, March 2019.

[18] Alex Krizhevsky, Ilya Sutskever, and Geoffrey E. Hinton. Imagenet classification with deep convolutional neural networks. In Proceedings of the 25th International Conference on Neural Information Processing Systems -Volume 1, NIPS’12, page 1097–1105, Red Hook, NY, USA, 2012. Curran Associates Inc.

[19] K.R. Kruthika Rajeswari, and H.D. Maheshappa. CBIR system using capsule networks and 3d CNN for alzheimer’s disease diagnosis. Informatics in Medicine Unlocked, 14:59–68, 2019.

[20] Xiu-Li Kuang, Xiao-Mei Zhao, Hai-Fang Xu, Yuan-Yuan Shi, Jin-Bo Deng, and Guo-Tao Sun. Spatio-temporal expression of a novel neuron-derived neurotrophic factor (NDNF) in mouse brains during development. BMC Neuroscience, 11(1):137, 2010.

[21] Woo-Young Lee, Seung-Min Park, and Kwee-Bo Sim. Optimal hyperparameter tuning of convolutional neural networks based on the parameter-setting-free harmony search algorithm. Optik, 172:359–367, nov 2018.

[22] Yutong Li, Ruoqing Zhu, Mike Yeh, and Annie Qu. Dermoscopic image classification with neural style transfer. Journal of Computational and Graphical Statistics, pages 1–14, may 2022.

[23] Olof Lindberg Mark … Eva Walterfang, and et al. Hippocampal shape analysis in alzheimer’s disease and frontotemporal lobar degeneration subtypes. Journal of Alzheimer’s Disease, 30(2):355–365, May 2012.

[24] Michelle F. Miranda, Hongtu Zhu, and Joseph G. Ibrahim. TPRM: Tensor partition regression models with applications in imaging biomarker detection. The Annals of Applied Statistics, 12(3):1422–1450, 2018.

[25] Alireza Nazarian, Anatoliy I. Yashin, and Alexander M. Kulminski. Genome-wide analysis of genetic predisposition to alzheimer’s disease and related sex disparities. Alzheimer’s Research & Therapy, 11(1), jan 2019.

[26] Chigozie Nwankpa, Winifred Ijomah, Anthony Gachagan, and Stephen Marshall. Activation functions: Comparison of trends in practice and research for deep learning. CoRR, abs/1811.03378, 2018.

[27] Yuqing Pan, Qing Mai, and Xin Zhang. Covariate-adjusted tensor classification in high dimensions. Journal of the American Statistical Association, 114(527):1305–1319, 2019.

[28] R C Petersen, P S Aisen, L A Beckett, M C Donohue, A C Gamst, D J Harvey, C R Jack, W J Jagust, L M Shaw, A W Toga, J Q Trojanowski, and M W Weiner. Alzheimer’s disease neuroimaging initiative (adni): clinical characterization. Neurology, 74:201–209, January 2010.

[29] Steven G. Potkin, Guia Guffanti, Anita Lakatos, Jessica A. Turner, Frithjof Kruggel, James H. Fallon, Andrew J. Saykin, Alessandro Orro, Sara Lupoli, Erika Salvi, Michael Weiner, and Fabio Macciardi. Hippocampal atrophy as a quantitative trait in a genome-wide association study identifying novel susceptibility genes for alzheimer’s disease. PLoS ONE, 4(8):e6501, aug 2009.

[30] Marco Tulio Ribeiro, Sameer Singh, and Carlos Guestrin. “why should i trust you?”: Explaining the predictions of any classifier. In Proceedings of the 22nd ACM SIGKDD International Conference on Knowledge Discovery and Data Mining, KDD ‘16, page 1135–1144, New York, NY, USA, 2016. Association for Computing Machinery.

[31] U. S. Sandau, M. Colino-Oliveira …M. J. Diogenes, A. M. Sebastiao, and D. Boison. Adenosine kinase deficiency in the brain results in maladaptive synaptic plasticity. Journal of Neuroscience, 36(48):12117–12128, nov 2016.

[32] Andrew J. Saykin, Li Shen, Tatiana M. Foroud, Steven G. Potkin, Shanker Swaminathan, Sungeun Kim, Shannon L. Risacher, Kwangsik Nho, Matthew J. Huentelman, David W. Craig, Paul M. Thompson, Jason L. Stein, Jason H. Moore, Lindsay A. Farrer, Robert C. Green, Lars Bertram, Clifford R. Jack, and Michael W. Weiner. Alzheimer’s disease neuroimaging initiative biomarkers as quantitative phenotypes: Genetics core aims, progress, and plans. Alzheimers & Dementia, 6(3):265–273, may 2010.

[33] Sharay E. Setti, Holly C. Hunsberger, and Miranda N. Reed. Alterations in hippocampal activity and alzheimer’s disease. Translational Issues in Psychological Science, 3(4):348–356, ec 2017.

[34] Li Shen, Sungeun Kim, Shannon L. Risacher, Kwangsik Nho, Shanker Swaminathan, John D. West, Tatiana Foroud, Nathan Pankratz, Jason H. Moore, Chantel D. Sloan, Matthew J. Huentelman, David W. Craig, Bryan M. DeChairo, Steven G. Potkin, Clifford R. Jack, Michael W. Weiner, and Andrew J. Saykin. Whole genome association study of brain-wide imaging phenotypes for identifying quantitative trait loci in MCI and AD: A study of the ADNI cohort. NeuroImage, 53(3):1051–1063, nov 2010.

[35] Ran Shi and Jian Kang. Thresholded multiscale gaussian processes with application to bayesian feature selection for massive neuroimaging data, 2015.

[36] Karl Sieg. Neurodevelopmental disorders associated with chromosome 15. Jefferson Journal of Psychiatry, 8(2), 1990.

[37] Karen Simonyan and Andrew Zisserman. Very deep convolutional networks for large-scale image recognition. CoRR, abs/1409.1556, 2015.

[38] Stephen M. Smith. Fast robust automated brain extraction. Human Brain Mapping, 17(3):143–155, nov 2002.

[39] Chun-I Sze. Role of WWOX WOX1 in alzheimer s disease pathology and in cell death signaling. Frontiers in Bioscience, E4(5):1951–1965, 2012.

[40] Xiwei Tang, Xuan Bi, and Annie Qu. Individualized multilayer tensor learning with an application in imaging analysis. Journal of the American Statistical Association, 115(530):836–851, 2020.

[41] Rui ting Hu, Qian Yu, Shao dan Zhou, Yi xin Yin, Rui guang Hu, Hai peng Lu, and Bang li Hu. Co-expression network analysis reveals novel genes underlying alzheimer’s disease pathogenesis. Frontiers in Aging Neuroscience, 12, nov 2020.

[42] Jade Xiaoqing Wang, Yimei Li, Xintong Li, and Zhao-Hua Lu. Alzheimer’s disease classification through imaging genetic data with IGnet. Frontiers in Neuroscience, 16, mar 2022.

[43] Mark W. Woolrich, Saad Jbabdi, Brian Patenaude, Michael Chappell, Salima Makni, Timothy Behrens, Christian Beckmann, Mark Jenkinson, and Stephen M. Smith. Bayesian analysis of neuroimaging data in FSL. NeuroImage, 45(1):S173–S186, mar 2009.

[44] Y. Zhang, M. Brady, and S. Smith. Segmentation of brain MR images through a hidden markov random field model and the expectation-maximization algorithm. IEEE Transactions on Medical Imaging, 20(1):45–57, 2001.

[45] Bingxin Zhao, Tengfei Li, Stephen M. Smith, Di Xiong, Xifeng Wang, Yue Yang, Tianyou Luo, Ziliang Zhu, Yue Shan, Nana Matoba, Quan Sun, Yuchen Yang, Mads E. Hauberg, Jaroslav Bendl, John F. Fullard, Panagiotis Roussos, Weili Lin, Yun Li, Jason L. Stein, and Hongtu Zhu. Common variants contribute to intrinsic human brain functional networks. jul 2020.

[46] Bingxin Zhao, Tengfei Li, Yue Yang, Xifeng Wang, Tianyou Luo, Yue Shan, Ziliang Zhu, D. Xiong, Mads E. Hauberg, Jaroslav Bendl, John F. Fullard, Panagiotis Roussos, Yun Li, Jason L. Stein, and Hongtu Zhu. Common genetic variation influencing human white matter microstructure. Science, 372(6548), jun 2021.

[47] Bingxin Zhao, Tianyou Luo, Tengfei Li, Yun Li, Jingwen Zhang, Yue Shan, Xifeng Wang, Liuqing Yang, Fan Zhou, Ziliang Zhu, Hongtu Zhu, and and. Genome-wide association analysis of 19,629 individuals identifies variants influencing regional brain volumes and refines their genetic co-architecture with cognitive and mental health traits. Nature Genetics, 51(11):1637–1644, nov 2019.

[48] Yize Zhao, Xiwen Zhao, Mansu Kim, Jingxuan Bao, and Li Shen. A novel bayesian semiparametric model for learning heritable imaging traits. In Medical Image Computing and Computer Assisted Intervention – MICCAI 2021, pages 678–687. Springer International Publishing, 2021.

