## Supplementary Table for "Deep Learning-based Feature Extraction with MRI Data in Neuroimaging Genetics for Alzheimer’s Disease"

**Supplementary Material**

Table 1: Accuracy Measures and associated SNPs and Genes

| Models | Training AUC | Testing AUC | SNPs | Genes |
|---|---|---|---|---|
| Whole-brain | 0.88 | 0.72 | 149 SNPs at 5 $\times 10^{-6}$ | APOE |
| Logistic Regression | 0.78 | 0.59 | GWAS Not Performed | |
| SVM | 0.79 | 0.61 | GWAS Not Performed | |
| ResNet50<br>AlexNet<br>VGG16 | 0.84-0.87 | 0.69-0.72 | GWAS Not Performed | |
| Augmentation with respect to whole-brain | 0.85 | 0.75 | 3 SNPs at 5 $\times 10^{-8}$<br>$rs$2075650, $rs$11580593<br>$rs$823955<br>38 SNPs at 5 $\times 10^{-6}$ | APOE |
| AD vs CN | 0.96 | 0.90 | 2 SNPs at 5 $\times 10^{-8}$<br>53 SNPs at 5 $\times 10^{-6}$ | ADCY8, ADK, APOE |
| Multibranch CNN (27 Models) | 0.86 | 0.76 | 8 SNPs at 5 $\times 10^{-8}$<br>87 SNPs at 5 $\times 10^{-6}$<br>$rs$1397645,$rs$10490381 | NDNF |
| Whole-image GM | 0.82 | 0.70 | No Significant genes were found | |
| Whole-image WM | 0.88 | 0.68 | No Significant genes were found | |
| Whole-image CSF | 0.87 | 0.54 | No Significant genes were found | |
| Multibranch GM | 0.84 | 0.74 | 35 SNPs at 5 $\times 10^{-6}$<br>$rs$173754,$rs$9257694 | APOE, OR14J1 |
| Multibranch WM | 0.82 | 0.70 | 35 SNPs at 5 $\times 10^{-6}$ | No Significant genes were found |
| Multibranch CSF | 0.77 | 0.66 | 21 SNPs at 5 $\times 10^{-6}$ | No Significant genes were found |

### Relevant Plots

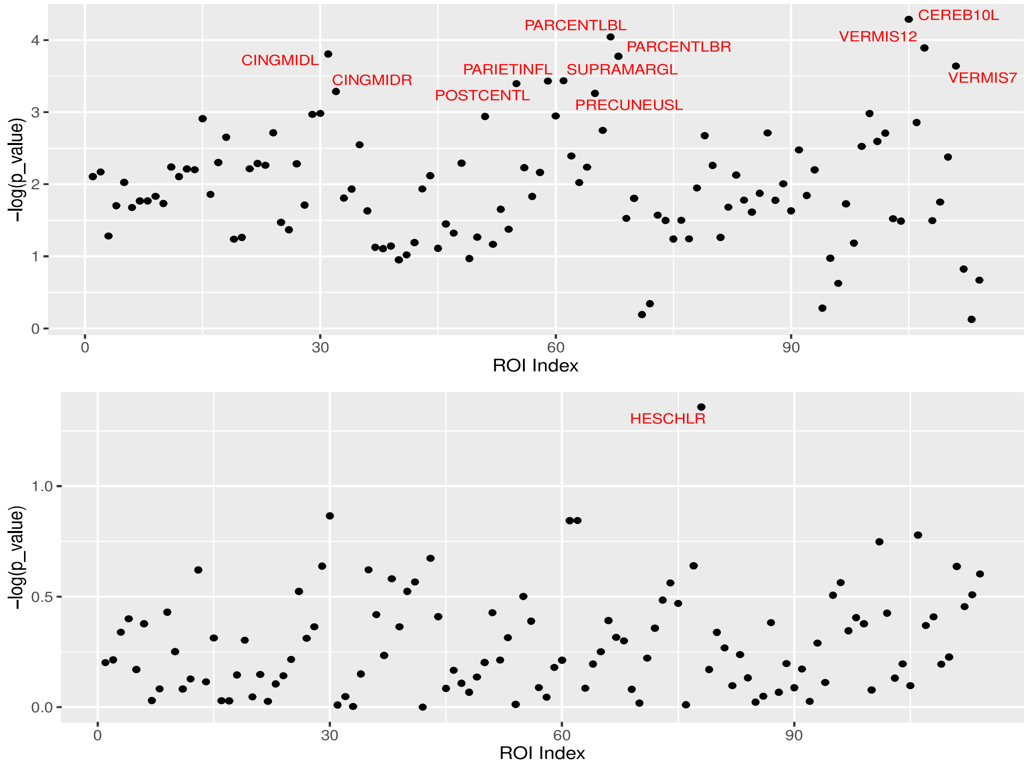


Figure 1: Not all the PCs are easy to interpret. The upper panel shows the plot of ROIs against the $-\log(p_value)$for PC1 and it is hard to pin-point one specific part of the ROIs. Similarly, in the lower panel we have HESCHL (transverse temporal gyri,right hemisphere) but the corresponding PC8 doesn't play a significant part in the identification of the important SNPs.
